# The effects of mindfulness on working memory: a systematic review and meta-analysis

**DOI:** 10.1101/2025.03.21.644687

**Authors:** Afsaneh Moradi, Maryam Ghorbani, Farzaneh Pouladi, Bridget Caldwell, Neil W. Bailey

## Abstract

**Objectives:** Mindfulness is a promising health intervention, showing potential effects on cognitive functions like memory. Despite evidence suggesting mindfulness improves working memory, inconsistencies in results and methodologies prevent definitive conclusions. This meta-analysis examines the effects of mindfulness interventions on working memory across clinical and healthy populations, and various age groups.

**Methods:** A systematic search for relevant English and Persian articles was conducted in WOS, Scopus, PsycINFO, and PubMed databases, along with nine meta-analyses up to February 2023. Included studies consisted of randomized controlled trials (RCTs), controlled trials (CTs), and single-group studies. Overall, 29 studies with 2076 participants aged 5–85 years were analyzed.

**Results:** Mindfulness interventions demonstrated a medium effect size on working memory: RCT two-group studies (Hedges g = 0.438, p < 0.001), CT two-group studies (Hedges g = 0.385, p < 0.005), and single-group studies (Hedges g = 0.583, p < 0.001). These findings confirm the effectiveness of mindfulness interventions in improving working memory.

**Conclusions:** Mindfulness interventions exhibit promising effects on working memory. However, further primary research, particularly rigorous RCTs, is needed to better understand their impacts on clinical versus healthy populations and across diverse age groups.

## Introduction

Mindfulness training has been widely incorporated into several intervention programs, such as Mindfulness-Based Stress Reduction (MBSR) and Mindfulness-Based Cognitive Therapy (MBCT). Other third-wave cognitive interventions also often include mindfulness practices, including Dialectical Behavior Therapy (DBT) and Acceptance and Commitment Therapy (ACT). Mindfulness was introduced to Western psychology by Kabat-Zinn in the early 1970s. According to his definition, mindfulness is “the ability to focus one’s attention on present-moment awareness while maintaining a non-judgmental attitude towards current experiences, including any emotions, thoughts, and sensations” (Kabat-Zinn, 1990). Similarly, in Buddhist philosophy, mindfulness is typically defined as the intentional attention to thoughts, emotions, and sensations as they arise moment by moment, accompanied by an open and accepting attitude (Millett et al., 2021a). While the concept of mindfulness reflects a specific quality of focused attention and mental awareness rather than the practice or use of a particular technique (Chambers et al., 2008), it is recognized that practice or training involving a continuous shift from an automatic mindset to one of attention and awareness can enhance a person’s dispositional mindfulness (Brown & Ryan, 2003).

Overall, mindfulness-based interventions have demonstrated positive effects on psychological functioning (Quach et al., 2016). Much of the early research on mindfulness focused on its ability to improve individual well-being through its beneficial effects on stress, anxiety, and depression (Lueke & Lueke, 2019). Studies have shown that mindfulness practices and mindfulness-based psychotherapy are successful in improving emotional and behavioral regulation, and reducing a wide range of psychological symptoms (Eisenbeck et al., 2018). Regardless of whether studies have examined clinical or healthy populations, mindfulness interventions have been shown to have effects involving reduced stress, distress, anxiety, and depression, and increased quality of life (Mirabito & Verhaeghen, 2023).

In addition to the effects for a person’s well-being, mindfulness training is understood to enhance specific neural and cognitive processes, including working memory (Morrison & Jha, 2015). Working memory is a higher-order cognitive process that is recognized to be one of the most important constructs in cognitive psychology (Scharfen et al., 2018). The concept of a working memory system was initially proposed by Baddeley and Hitch (1974) as a dynamic multi-system short-term memory (Levi & Rosenstreich, 2019). Working memory is now recognized as a cognitive function that provides temporary storage and manipulation of information necessary for complex cognitive tasks (Melby-Lervåg & Hulme, 2013). This memory allows us to retain and manipulate goal-relevant information over short intervals, without being distracted by irrelevant information, which allows adaptive behavior (Jha et al., 2020). Since the maintenance of a person’s goals in working memory is required to maintain attention over time, including the goal to maintain mindful awareness during mindfulness training, researchers have argued that mindfulness training is could also be considered to be a form of working memory practice, and as a result, might be associated with enhancement of working memory function (Mrazek et al., 2013). For example, when an individual engages in a mindful breathing exercise, working memory processes must support attention to the breath over time and protect against unrelated thoughts or external stimuli (Goyal et al., 2014; Zeidan et al., 2010).

In addition to the putative “working memory practice effects” of mindfulness, the practice of mindfulness has also been suggested to produce improved attention control, which may enable an individual to use working memory more efficiently by limiting the processing of irrelevant information (Morrison & Jha, 2015). Mindfulness might also indirectly reduce memory problems by reducing the perceived stress that can adversely affect working memory function (Brisbon & Lachman, 2017). Indeed, studies show that mind wandering, or having scattered thoughts during ongoing activities, is associated with lower working memory function (Jha et al., 2019).

Given these rationales for the suggestion that mindfulness practice might improve memory, a number of studies have examined the impact of mindfulness on various types of memory, including working memory. Many of these studies have shown that mindfulness does enhance memory (Lueke & Lueke, 2019). These effects have been noted across different measures of memory (for example verbal learning and working memory capacity: (Dubert et al., 2016; Lueke & Lueke, 2019), and across healthy (Brown et al., 2016; Shemesh et al., 2023) and clinical (Bachmann et al., 2018; Isham et al., 2020) samples. In a meta-analysis examining the impact of mindfulness on executive control conducted on adults, a significant effect with a medium effect size was found for the effects of mindfulness on working memory (Cásedas et al., 2020). Similarly, a meta-analysis conducted to investigate the effects of mindfulness-based programs across a range of cognitive functions indicated that mindfulness had the greatest impact on working memory compared to other executive function indicators (Whitfield et al., 2022).

However, reviews have indicated that the effects of mindfulness might not be consistent across all memory functions. In a meta-analysis on the impact of mindfulness, it was found that while mindfulness had small to moderate effects on working memory accuracy, it did not affect episodic memory (Zainal & Newman, 2024). The effects of mindfulness on working memory are also not consistently reported (for example, (Flook et al., 2024; Lovette et al., 2022). Meta-analysis by Yakobi et al. (2021) reported significant effects of mindfulness interventions on attention and executive control, but no effect on working memory (Yakobi et al., 2021). Similarly, in a meta-analysis conducted with group-based mindfulness interventions, although a very small effect of mindfulness on working memory was observed, the overall results indicated that the impact of mindfulness interventions on executive functions is not particularly strong (Millett et al., 2021a). Overall, despite some evidence suggesting that mindfulness meditation may improve cognition, inconsistencies across studies may have limited definitive conclusions (Gill et al., 2020). While there is a reasonable volume of research on the effects of mindfulness on working memory, meta-analyses in this area are limited. Most of the existing meta-analyses have examined the effectiveness of mindfulness in non-clinical groups with a focus on assessing the effects of mindfulness on a range of cognitive functions, with working memory being only part of the set (Cásedas et al., 2020; Whitfield et al., 2022; Yakobi et al., 2021). Other approaches have examined the effects of mindfulness on executive functions in individuals over 18 years old, excluding children and adolescents (Chiesa et al., 2011; Im et al., 2021; Millett et al., 2021b). In contrast, the present study specifically examined the effectiveness of mindfulness on working memory, including both clinical and non-clinical groups and all age ranges. We note that a recent review has examined the impact of mindfulness on various executive functions (Zainal & Newman, 2024). However, despite similarities in some inclusion criteria (for example, using no age restrictions and including both clinical and healthy populations), there are differences between their analysis and the present study. Zainal & Newman ’s meta-analysis included only randomized controlled trials (RCTs), whereas the present study included both RCTs and nonrandomized controlled trials (CTs). The meta-analysis by Zeinal & Newman (2024) also included fewer studies that focused solely on WM than the present study. Therefore, the aim of the present study was to determine the effect sizes of mindfulness on working memory through a comprehensive meta-analysis including single-group studies and two-group studies, and including both RCT and CT methods, both clinical and healthy populations, and all age ranges.

## Method

### Literature Search

This study is a meta-analysis based on the PRISMA guidelines (Moher et al., 2009) aimed at examining the impact of mindfulness on working memory. A systematic search of English and Persian articles was conducted in databases including WOS, Scopus, PsycINFO, and PubMed up to February 2023. The keywords used were “working memory,” “mindfulness,” “MBSR,” “meditation,” “MBCT,” and various combinations of these terms. The search strategy was as follows: (TITLE (("mindfulness” OR “meditation” OR “mbsr” OR “mbct") AND “working memory") OR KEY (("mindfulness” OR “meditation” OR “mbsr” OR “mbct") AND “working memory")) AND (LIMIT-TO(DOCTYPE, “ar") OR LIMIT-TO(DOCTYPE, “re")) AND (LIMIT- TO(LANGUAGE, “English”) OR LIMIT-TO (LANGUAGE, “Persian”)) AND (LIMIT-TO(SRCTYPE, “j")) AND (LIMIT-TO(PUBSTAGE, “final")). Additionally, articles related to the impact of mindfulness interventions on working memory in the reference lists of 9 previous meta-analyses (Table 1) were manually searched and reviewed. In total, 359 studies were identified. In the initial screening phase, the titles and abstracts of the studies were reviewed to check for duplicates, article type (article), and the presence of an abstract. In the next phase, articles were evaluated based on inclusion and exclusion criteria. The studies selected for analysis were then screened based on the full text to confirm eligibility. Each study was independently screened by two reviewers, and discrepancies were resolved through discussion. Finally, 29 studies were selected for the final analysis. For the complete screening process, refer to the flowchart (Moher et al., 2009) in Figure 1.

**Fig 1.**
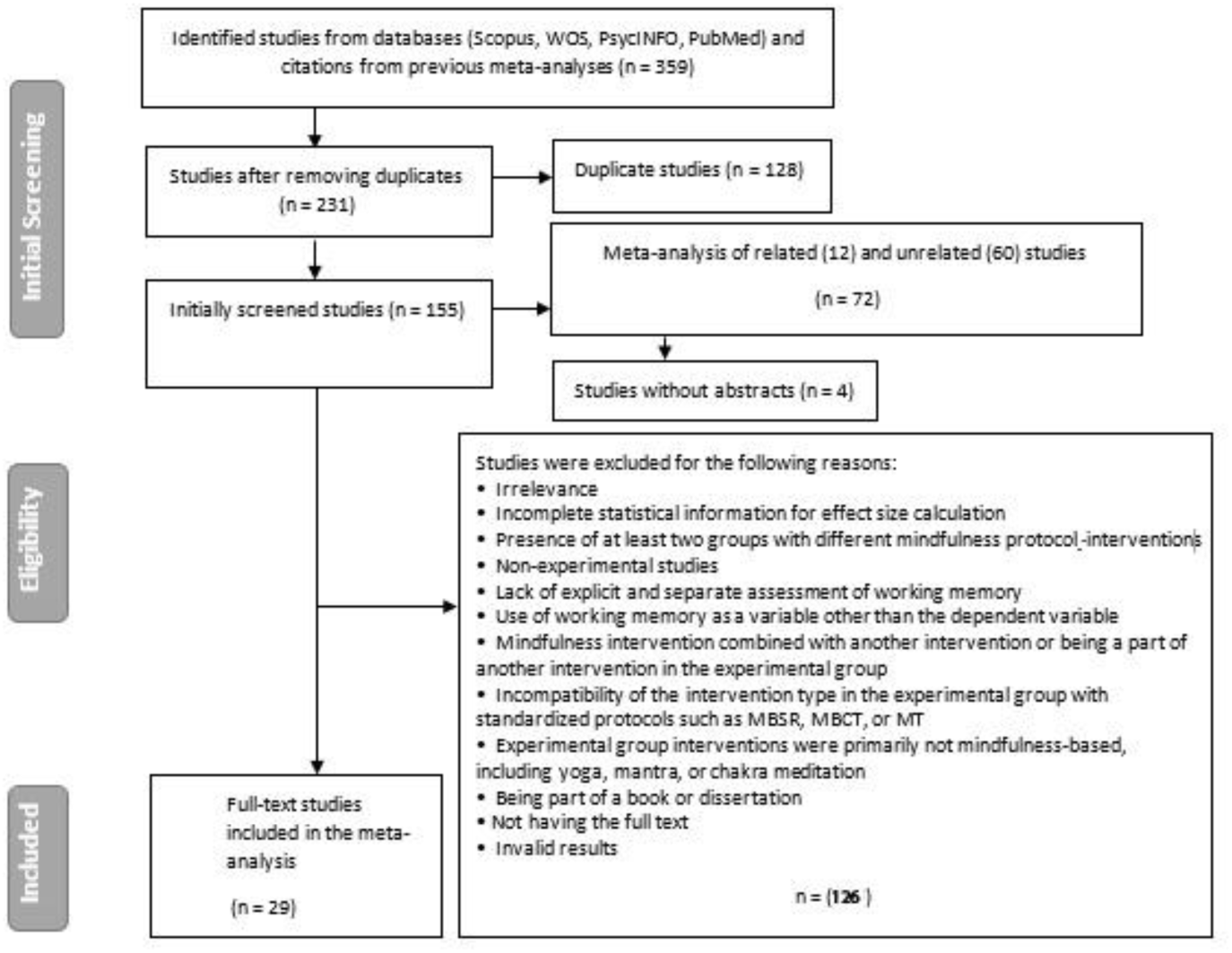
Systematic review flow diagram. Adapted from “Preferred Reporting Items for Systematic Reviews and Meta Analyses: The PRISMA Statement (Moher et al., 2009).

**Table 1.**
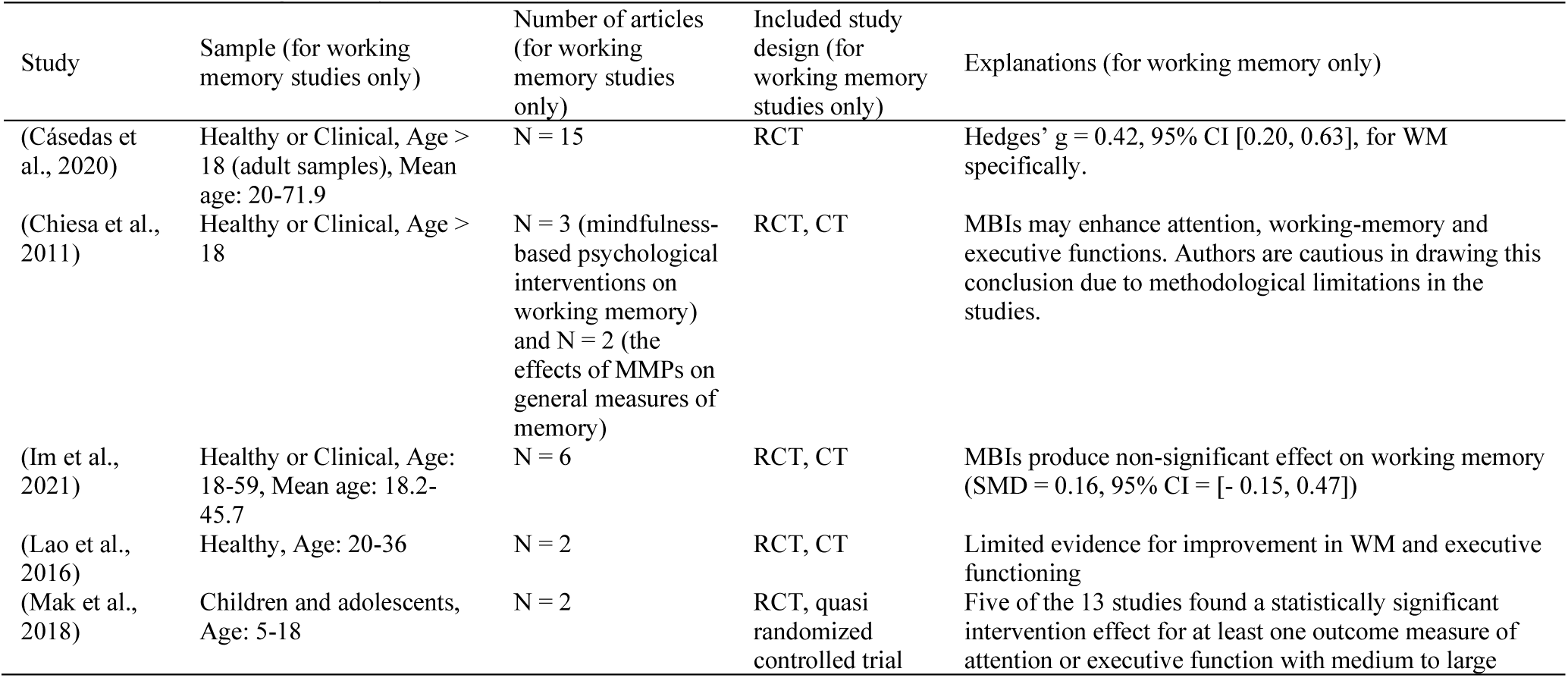
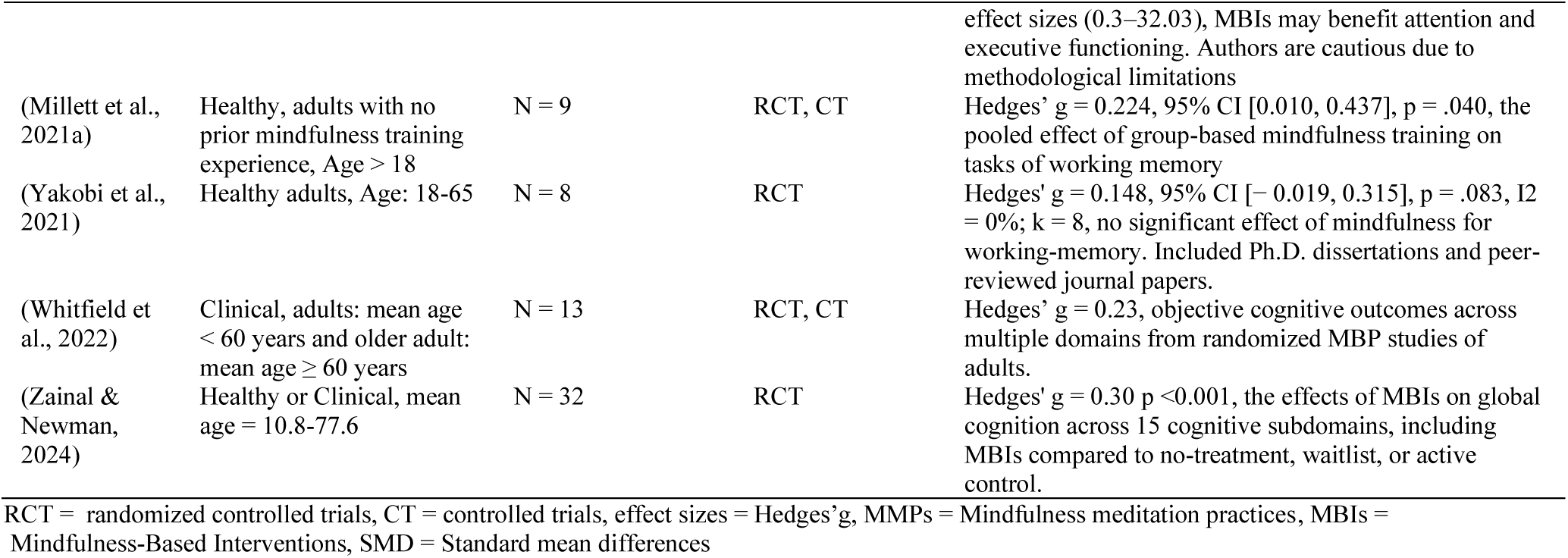
Findings from previous systematic reviews and meta-analyses with more than one study on the effect of mindfulness interventions on working memory (n = 9)

### Study Selection

Studies that included mindfulness interventions and examined working memory were selected without age restrictions for participants. Studies focusing on yoga, mantra, or chakra meditation were excluded. Additionally, studies that did not include mindfulness interventions (e.g., studies examining only the relationship between error processing metrics and trait mindfulness) were also excluded from this analysis. Mindfulness-Based Interventions (MBIs) were required to include standardized protocols where mindfulness meditation is practiced, such as MBCT or MBSR. These protocols were also included if they were presented in various forms with minor deviations (e.g., using smartphone apps or audio recordings rather than being taught face to face) or presented with different implementation times. Furthermore, mindfulness practice was required to be the main component of the intervention and was required to not be combined with another activity, or to be a subsidiary part of another intervention. We included RCTs, CTs, and single-group studies (where a single-group received a mindfulness intervention and their baseline scores were compared with their post-intervention scores). In two-group studies, MBIs were compared with inactive or active control groups, and in single-group studies, changes in participants’ working memory before and after MBIs were examined. Full texts of studies that passed abstract screening were selected for inclusion in our meta-analysis based on the inclusion criteria specified in Table 2.

**Table 2.**
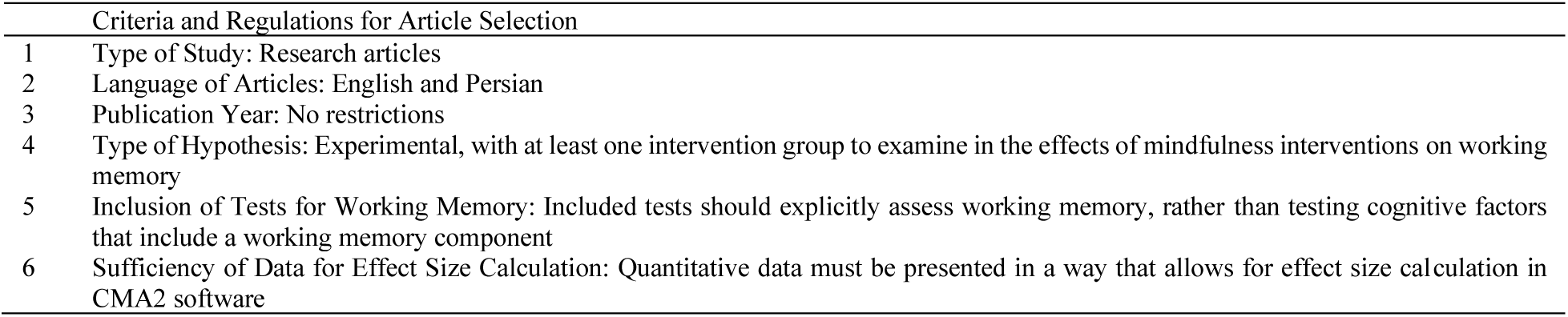
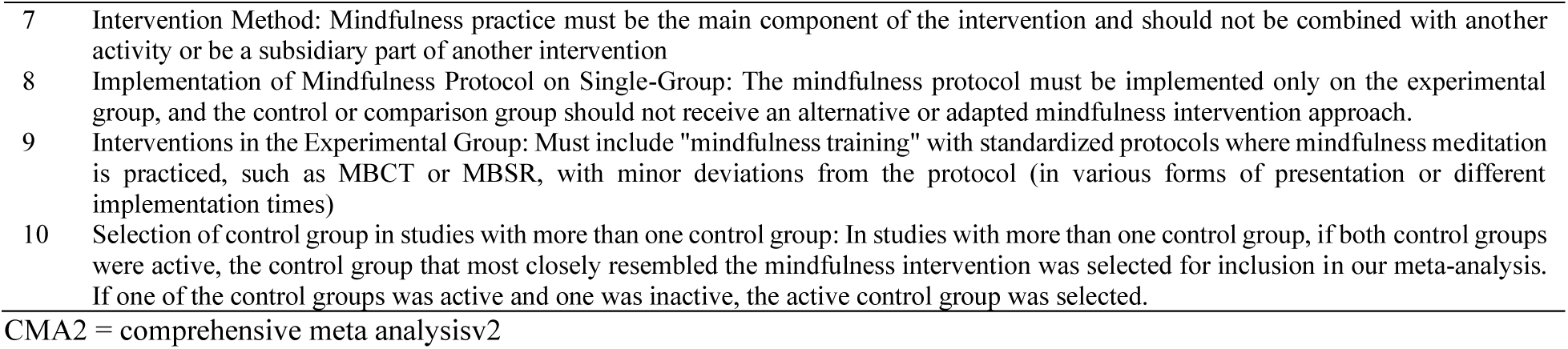
Selection Articles.

|  | Criteria and Regulations for Article Selection |
| --- | --- |
| 1 | Type of Study: Research articles |
| 2 | Language of Articles: English and Persian |
| 3 | Publication Year: No restrictions |
| 4 | Type of Hypothesis: Experimental, with at least one intervention group to examine in the effects of mindfulness interventions on working memory |
| 5 | Inclusion of Tests for Working Memory: Included tests should explicitly assess working memory, rather than testing cognitive factors that include a working memory component |
| 6 | Sufficiency of Data for Effect Size Calculation: Quantitative data must be presented in a way that allows for effect size calculation in CMA2 software |
| 7 | Intervention Method: Mindfulness practice must be the main component of the intervention and should not be combined with another activity or be a subsidiary part of another intervention |
| 8 | Implementation of Mindfulness Protocol on Single-Group: The mindfulness protocol must be implemented only on the experimental group, and the control or comparison group should not receive an alternative or adapted mindfulness intervention approach. |
| 9 | Interventions in the Experimental Group: Must include "mindfulness training" with standardized protocols where mindfulness meditation is practiced, such as MBCT or MBSR, with minor deviations from the protocol (in various forms of presentation or different implementation times) |
| 10 | Selection of control group in studies with more than one control group: In studies with more than one control group, if both control groups were active, the control group that most closely resembled the mindfulness intervention was selected for inclusion in our meta-analysis. If one of the control groups was active and one was inactive, the active control group was selected. |

CMA2 = comprehensive meta analysisv2

### Data Extraction

The extracted information from the articles included: general study information (author and year of publication), participant demographics (age and gender distribution), clinical characteristics of the subjects (presence of psychiatric disorders or healthy individuals), research design (RCT, CT, single group), and methodological features (sample size in experimental and control groups, sampling method, type of experimental design). Additionally, experimental group interventions details (type of mindfulness intervention, number of sessions, duration of intervention), features of control groups (active or inactive, name of intervention in active control), working memory tool characteristics (tools for measuring working memory), and statistical features (means, standard deviations, effect sizes, statistical significance) were also reviewed. Statistical analysis for each study was conducted to create a common metric (the effect size). The effect size was calculated using Hedges’ g as an estimate of effect size for each study to provide an unbiased estimate of sample size (Hedges, 1981). Hedges’ g also corrects for bias in small sample sizes (Lakens, 2013). Two researchers extracted the information and data from the literature to ensure their accuracy, with any discrepancies resolved via discussion and consultation with a third researcher if necessary.

### Study Quality

We assessed study quality by using the adapted version of the Delphi criteria (Verhagen et al., 1998) applied by Zainal and Newman (2024). This assessment criteria scored studies from 0 to 6 (with 6 reflecting the highest quality of study) using the following individual criterion: 1) participants were randomly allocated to either the intervention or the control group, 2) the groups were demographically similar, 3) the study reported the participant eligibility criteria, 4) the experimenters who administered the working memory assessments were blinded to participant group allocation, 5) statistics reported were sufficient to enable effect size calculation for inclusion in the meta-analysis, 6) an intention-to-treat analysis approach was used. Quality assessments were performed independently by two researchers (NWB and BC). Where conflicts in scoring were present, the allocated score was discussed with reference to the details reported in the manuscript, and conflicts were resolved by consensus between both of these researchers.

### Statistical Analysis

All statistical analyses were conducted using Comprehensive Meta-Analysis (CMA) Version 2. Effect sizes for each comparison of working memory performance was calculated using Hedges’ g, a weighted sample size which corrects for small sample bias (Hedges and Olkin, 1985). This effect size can be small (<0.2), medium (between 0.2 and 0.5), or large (>0.8) (Hedges & Olkin, 2014).

In this study, single-group studies, two-group RCTs, and CTs were separated and analyses were performed independently for each type of study. Hedges’ g effect size was calculated for each study based on post-intervention measurements for studies with two groups (mindfulness intervention and control) using a range of different methods, depending on the statistics reported by the original authors. For two-group studies, the “unmatched groups, post-only” method was used (independent groups (means, SD’s) was the preferred method, used for 18 studies (15 RCT studies and 3 CT studies), and the independent groups (Sample size, p) method was used for 5 RCT studies). Where the statistics reported did not enable the “unmatched groups, post-only” method to be used, the “unmatched groups, pre and post data” method was used (via the F for differences in change method used for 4 studies (3 RCT studies and 1 CT study)). Given that pre- and post-treatment correlations were not available in the “F for difference in change” method, we adopted a conservative estimate (r = 0.50) that reflects common test-to-retest correlations of cognitive function tasks reported in the literature to date (Dai et al., 2019; Hedge et al., 2018; Zainal & Newman, 2024).

For studies that included only a single-group (which examined pre- to post changes in working memory from a mindfulness intervention) or reported statistics that only enabled effect size calculation for single-group, the “single-group (pre, post) and matched groups” method was used (Paired groups (N, t-value)). A positive Hedges effect size indicates larger values in post-intervention tests compared to pre-intervention tests in single-group studies and greater effectiveness of the experimental group intervention compared to the control group in two-group studies (Bornstein et al., 2021; Hedges & Olkin, 2014).

After extracting these values from each study, a univariate meta-analysis was conducted separately for RCTs, CTs, and single-group studies to evaluate the statistical significance of the effect size across all included studies. The analysis accounted for heterogeneity by employing both fixed-effects and random-effects models.

Homogeneity tests were performed to determine whether the studies had similar distributions and to choose the fixed effects model as the main model or, in the case of heterogeneity, to select the random effects model. The heterogeneity of effect sizes was examined using Cochran’s Q test (Q Value) and I-squared. Cochran’s Q test indicates whether heterogeneity exists in the meta-analysis (with a significant P-value in this test indicating the presence of heterogeneity), and the I-squared test shows the percentage of variance among the included studies attributable to heterogeneity. A value of zero indicates homogeneity, while values of 25, 50, and 75 indicate low, medium, and high levels of heterogeneity, respectively (Higgins et al., 2003; Higgins & Thompson, 2002; Huedo-Medina et al., 2006). Finally, the included studies were assessed for various types of bias (Orwin, 1983; Rosenthal, 1979). One of the most significant biases in meta-analyses is publication bias, meaning that a study may never be published and remain in personal archives, which could alter meta-analytical results if published. To address the potential for publication bias in this research, the funnel plot and Duval and Tweedie’s trim and fill method was used (Duval & Tweedie, 2000). The funnel plot assumes that small effect sizes are prone to error and uses that assumption to assess the potential that some missing studies could have been included in the meta-analysis. A symmetrical distribution of articles around the effect size in the funnel plot indicates no publication bias, while an accumulation of studies on one side indicates publication bias (De Grijs et al., 2014; Nakagawa et al., 2022). The number of missing studies was estimated using Duval and Tweedie’s trim and fill method (Podina et al., 2016), and the robustness of our results against potential publication bias was assessed using the Fail-Safe N test for quality assessment. The Fail-Safe N analysis is used to evaluate the stability of results and the potential impact of publication bias. This analysis shows how many studies with non-significant results would need to be added to the meta-analysis to change the overall results from statistically significant to not significant. When a meta-analysis examines studies with large sample sizes, the Fail-Safe N value will be high, indicating a greater stability of results against publication bias and that many null result studies would be needed to change the meta-analysis results.

### Moderator Analyses

To explore the factors contributing to variations in the effect sizes of mindfulness-based interventions (MBIs), six potential moderators were assessed: (i) the type of control group, distinguishing between active and inactive groups; (ii) clinical status, comparing clinical populations with healthy individuals; (iii) the delivery method (FF: face-to-face sessions involving direct instructor-participant interaction versus ON: online formats excluding in-person engagement); (iv) duration of the intervention (DI: short ≤ 4 weeks or extended > 4 weeks); (v) the number of sessions (NS: low ≤ 4 sessions or high > 4 sessions); and (vi) the length of sessions (LS: brief ≤ 60 minutes or extended > 60 minutes). This analysis aimed to identify moderators that influence MBI outcomes.

## Results

Based on the search strategy, systematic literature review of the databases mentioned in the study selection section, and the review of articles from previous meta-analyses related to the aims of the current research (Table 1), our search resulted in 359 articles. A systematic filtering process was applied to these studies (see Figure 1). Initially, after removing duplicates, 231 studies remained. Following the initial screening, 155 articles were retained for inclusion assessment. Based on the inclusion and exclusion criteria, finally, the researcher summarized 29 full-text studies for statistical analysis.

### Study Characteristics

Table 3 contains the main characteristics of the studies included in our meta-analysis. The total number of participants included in our meta-analysis was 2076 (47 participants in single-group studies, 1818 participants in RCT two-group studies, 211 in CT two-group studies). Within the RCT two-group studies, 928 participants were in the mindfulness groups and 890 participants were in the control groups. Within the CT two-group studies, 111 participants were in the mindfulness groups and 100 participants were in the control groups. A total of 631 participants were in the RCT active control groups and 259 were in the inactive control groups, 80 participants were in the CT active control groups and 20 were in the inactive control groups. The age of participants ranged from 5 to 85 years. Additionally, 3 studies involved three groups, but the participants in the third group were not included in the current meta-analysis in order to avoid including statistics from the same group multiple times (Axelsen et al., 2022; Vieth & von Stockhausen, 2022).

**Table 3.**
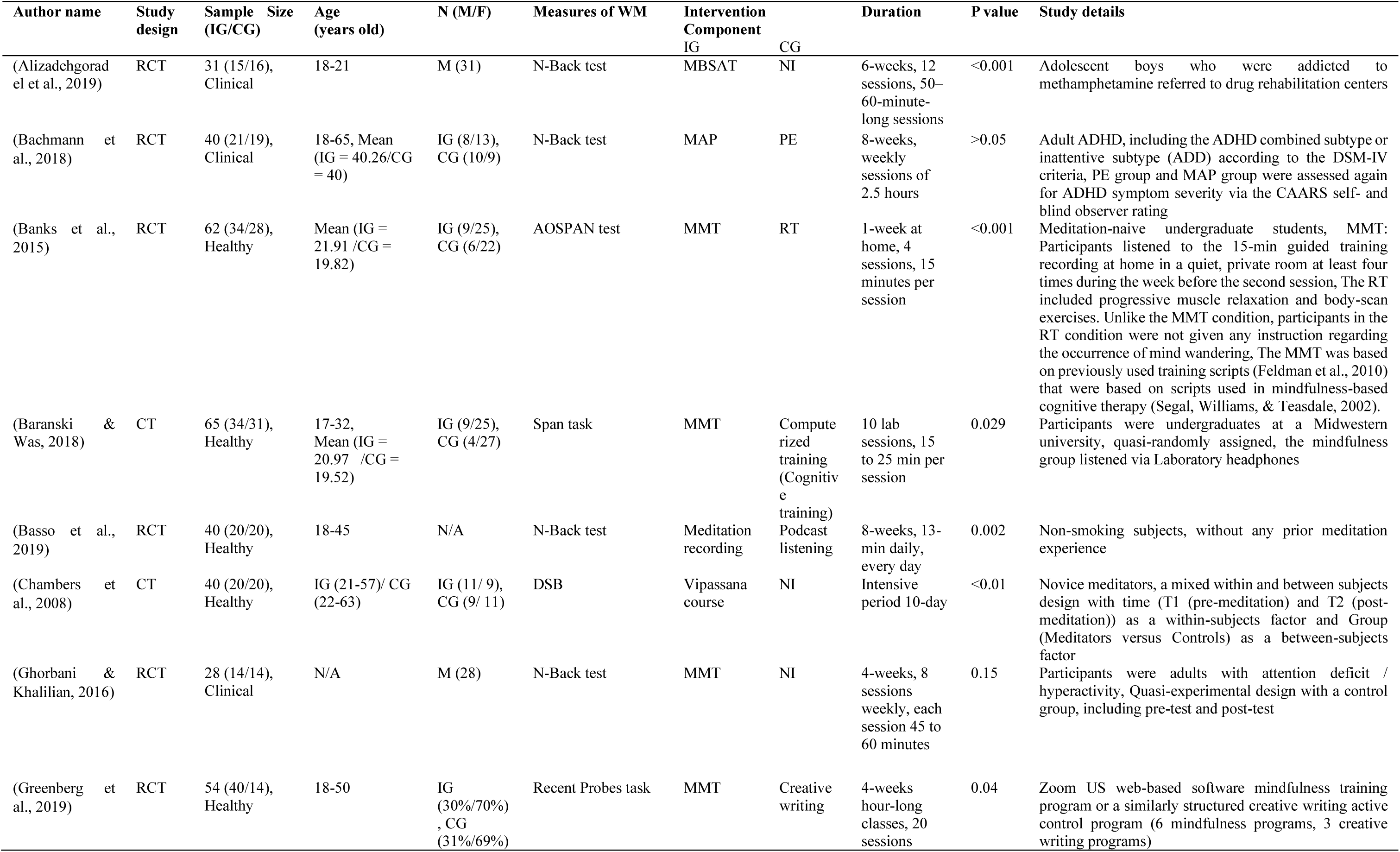
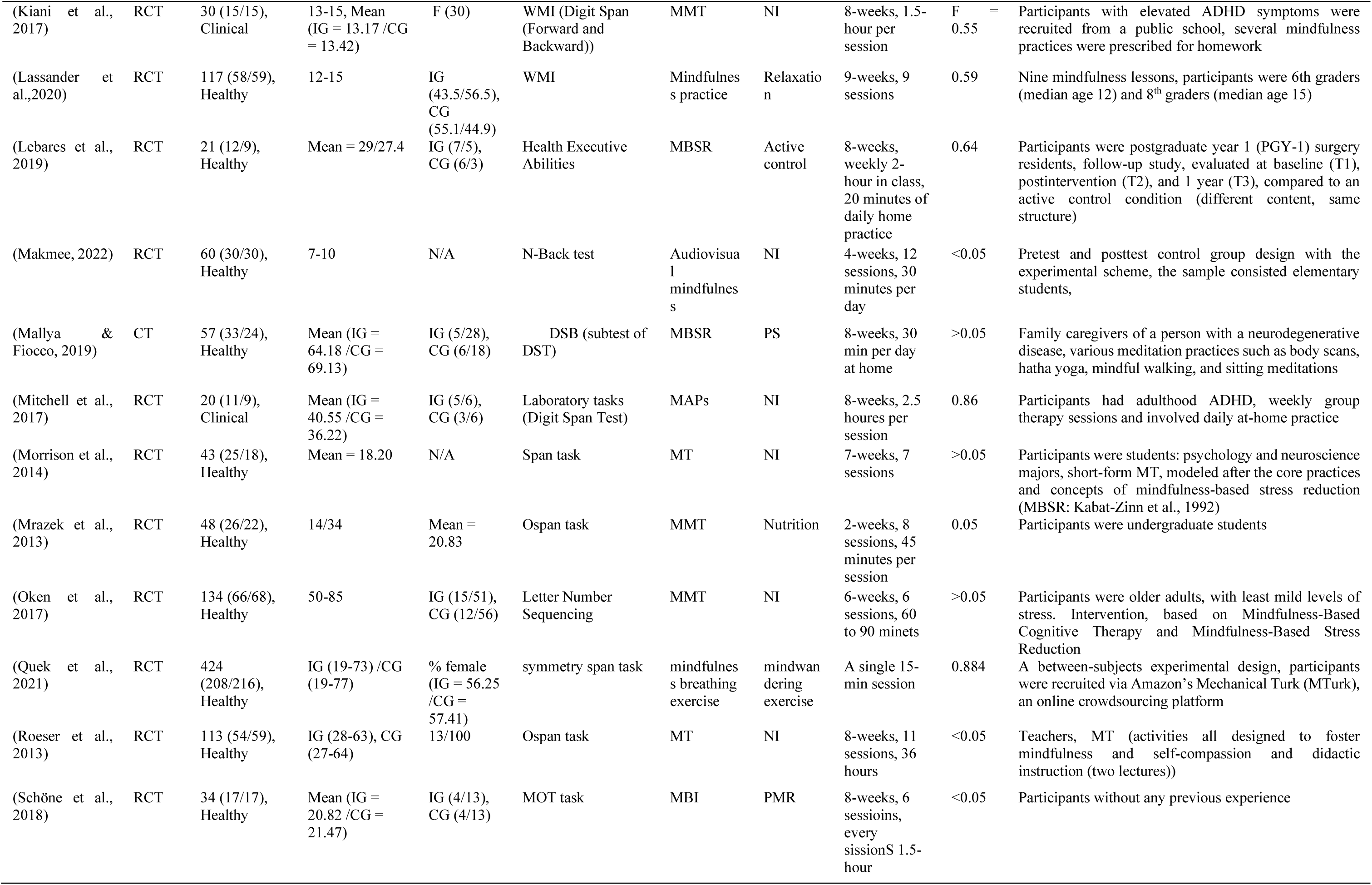
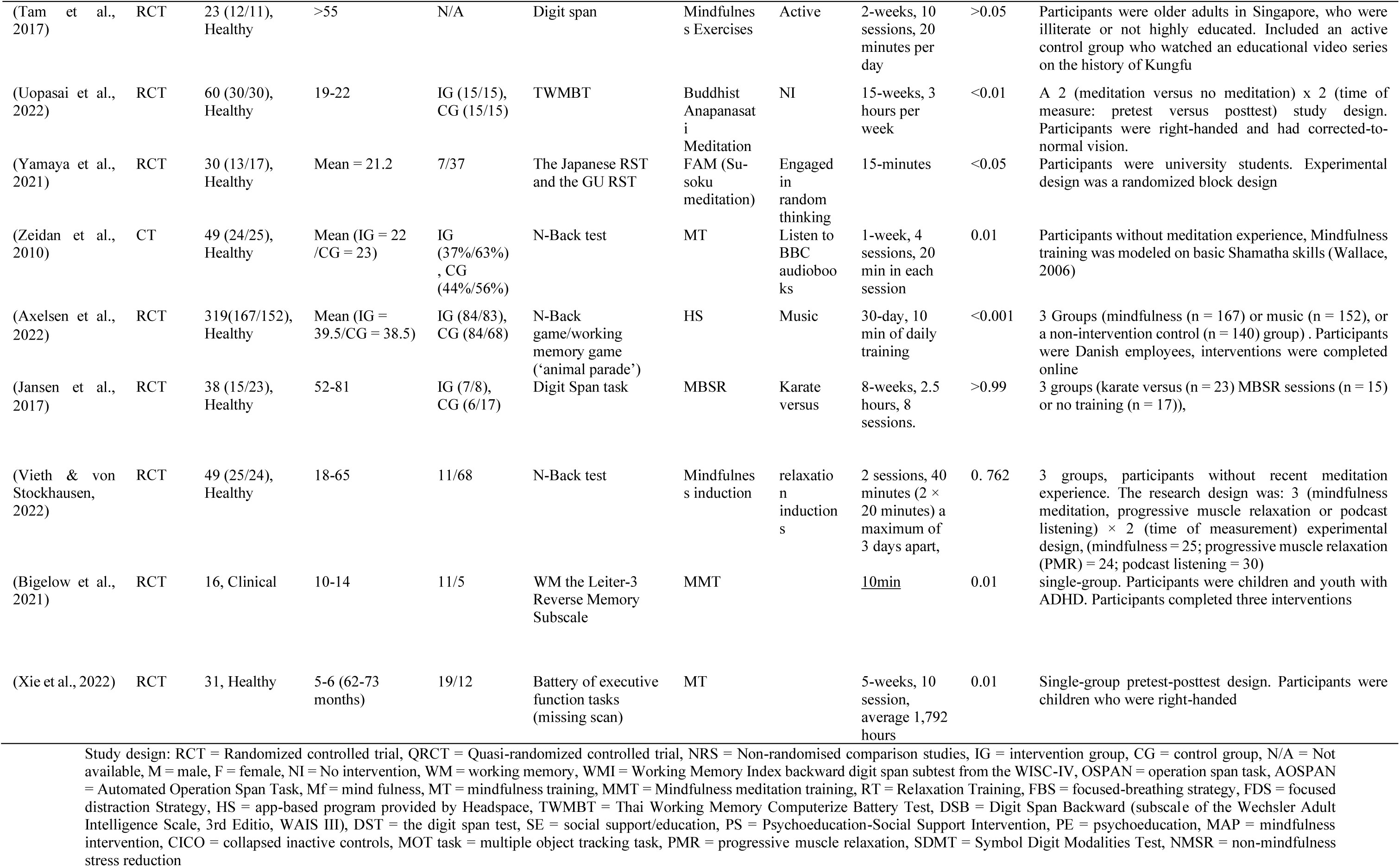
Characteristics of studies included in the meta-analyses (n = 29).

### Meta-analysis

#### Statistics

In Table 4, Table 5, and Table 6, the effect size of each study is presented along with the 95% confidence interval, lower and upper bounds, z-value, and p-value, indicating the significance of the effect size. Each study’s results were converted to Hedges’ g effect size using the appropriate formula. None of the random and fixed effect sizes showed significant differences from each other (all p-value < 0.001), and all models indicate a significant impact of mindfulness on working memory. Therefore, we proceed to examine the homogeneity and the consistency across the studies. For the studies that used a RCT two-group study design, the values of the fixed and random effect sizes were Hedges’ g = 0.367 and Hedges’ g = 0.438. For CT two-group study design, the values of the fixed and random effect sizes were Hedges’ g = 0.385 and Hedges’ g = 0.390 respectively. For the studies using a single-group design, the values of the fixed and random effect sizes were both Hedges’ g = 0.583 (although we note that only two studies were included in this analysis). Given the fixed effect size, random effect size and p-value in Tables 4, 5 and 6, the research hypothesis of a positive effect of mindfulness on working memory was confirmed at the 99% confidence level. Forest plots were drawn based on the effect size values for each study type, as well as the lower limit, and the upper limit bounds to assess the overall effects across studies (Figures 2 to 4).

**Fig 2.**
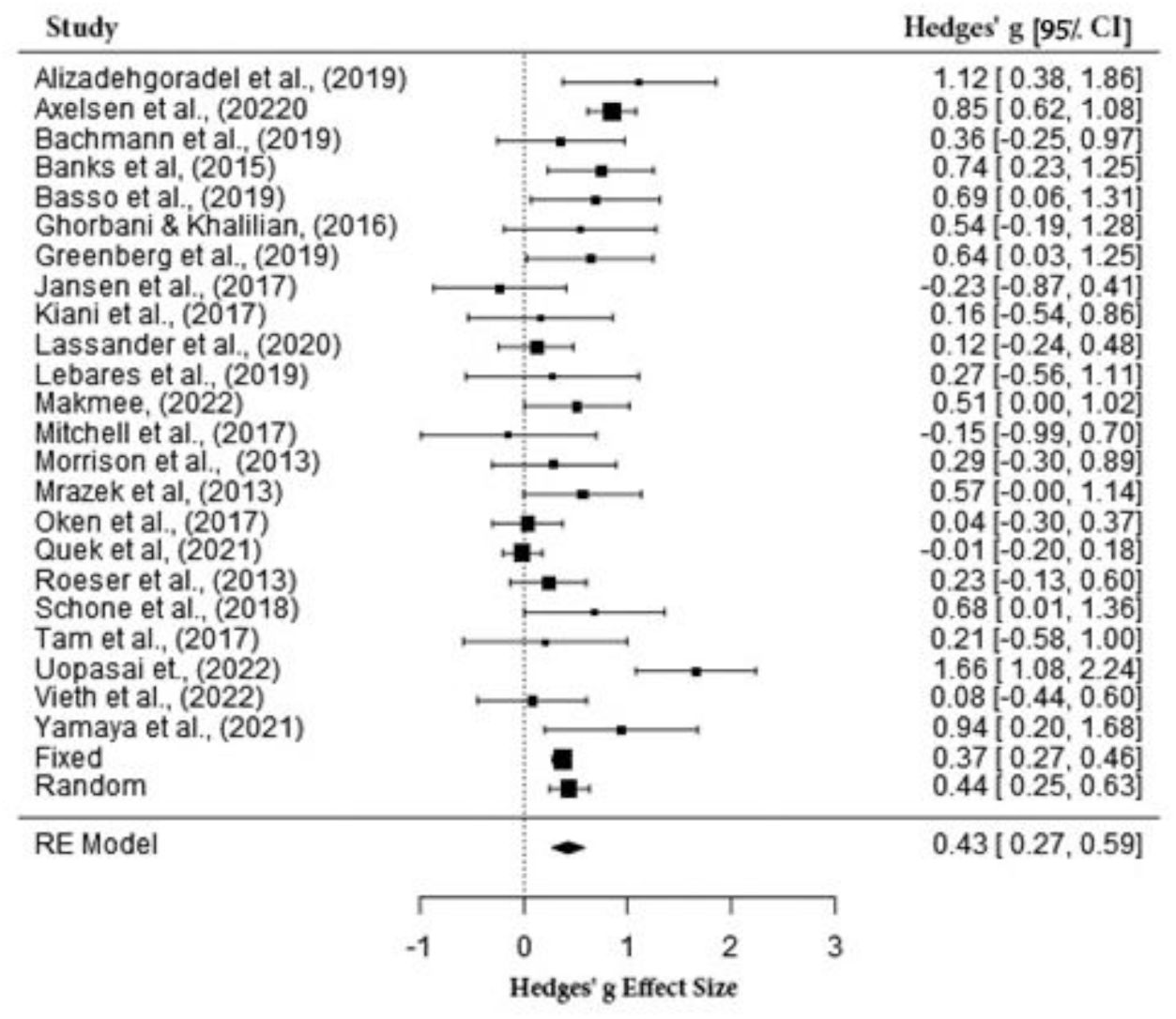
Forest plot of RCT 2-group studies included in meta-analysis for working memory

**Fig 3.**
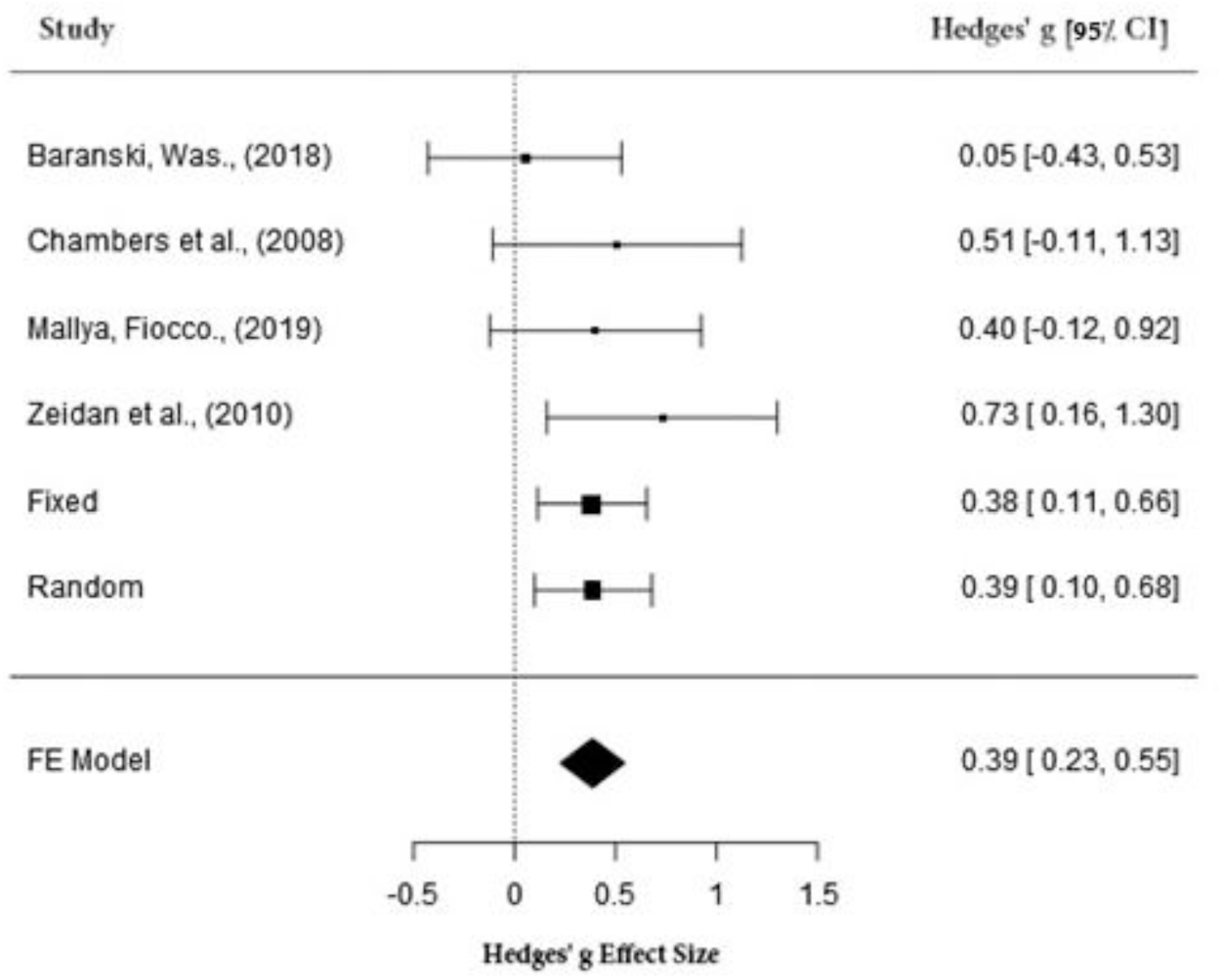
Forest plot of CT 2-group studies included in meta-analysis for working memory

**Fig 4.**
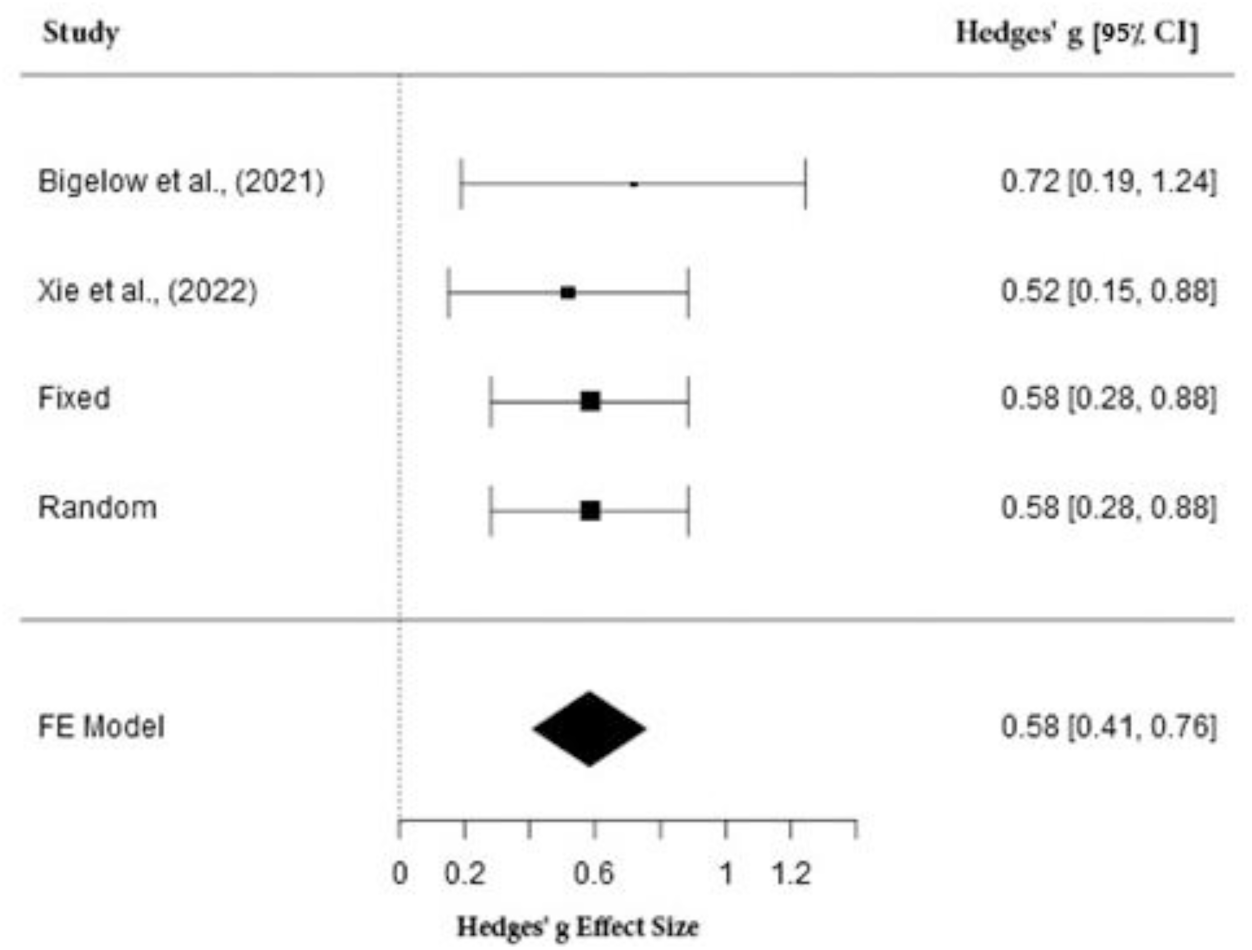
Forest plot of single-group studies included in meta-analysis for working memory

**Table 4.** Summary effect sizes of the RCT two-group studies (n = 23)

| Study name | Effect size | Lower limit | Upper limit | Z value | P value |
| --- | --- | --- | --- | --- | --- |
| Alizadehgoradel et al., (2019) | 1.118 | 0.378 | 1.859 | 2.961 | 0.003* |
| Axelsen et al., (2022) | 0.850 | 0.621 | 1.078 | 7.274 | 0.000** |
| Bachmann et al., (2019) | 0.359 | -0.254 | 0.972 | 1.147 | 0.251 |
| Banks et al., (2015) | 0.742 | 0.231 | 1.253 | 2.847 | 0.004* |
| Basso et al., (2019) | 0.689 | 0.063 | 1.315 | 2.158 | 0.031* |
| Ghorbani & Khalilian, (2016) | 0.544 | -0.189 | 1.278 | 1.455 | 0.146 |
| Greenberg et al., (2019) | 0.640 | 0.028 | 1.252 | 2.049 | 0.040* |
| Jansen et al., (2017) | -0.229 | -0.868 | 0.410 | -0.703 | 0.482 |
| Kiani et al., (2017) | 0.159 | -0.539 | 0.856 | 0.446 | 0.656 |
| Lassander et al., (2020) | 0.119 | -0.241 | 0.480 | 0.649 | 0.517 |
| Lebares et al., (2019) | 0.274 | -0.560 | 1.108 | 0.644 | 0.520 |
| Makmee, (2022) | 0.512 | 0.005 | 1.020 | 1.978 | 0.048* |
| Mitchell et al., (2017) | -0.149 | -0.994 | 0.696 | -0.345 | 0.730 |
| Morrison et al., (2013) | 0.295 | -0.303 | 0.893 | 0.968 | 0.333 |
| Mrazek et al., (2013) | 0.567 | -0.003 | 1.136 | 1.949 | 0.051 |
| Oken et al., (2017) | 0.037 | -0.300 | 0.374 | 0.214 | 0.830 |
| Quek et al., (2021) | -0.014 | -0.204 | 0.176 | -0.141 | 0.888 |
| Roeser et al., (2013) | 0.234 | -0.134 | 0.602 | 1.245 | 0.213 |
| Schoon et al., (2018) | 0.685 | 0.009 | 1.362 | 1.986 | 0.047* |
| Tam et al., (2017) | 0.209 | -0.581 | 1.000 | 0.519 | 0.604 |
| Uopasai et., (2022) | 1.661 | 1.080 | 2.242 | 5.602 | 0.000** |
| Vieth et al., (2022) | 0.081 | -0.442 | 0.605 | 0.304 | 0.761 |
| Yamaya et al., (2021) | 0.940 | 0.198 | 1.681 | 2.483 | 0.013* |
| Fixed | 0.367 | 0.274 | 0.460 | 7.734 | <0.001** |
| Random | 0.438 | 0.246 | 0.630 | 4.464 | <0.001** |

\*p<0.05 \*\*p<0.001

**Table 5.** Summary effect sizes of the CT two-group studies (n = 4)

| Study name | Effect size | Lower limit | Upper limit | Z value | P value |
| --- | --- | --- | --- | --- | --- |
| Baranski, Was., (2018) | 0.052 | -0.429 | 0.533 | 0.212 | 0.832 |
| Chambers et al., (2008) | 0.508 | -0.110 | 1.125 | 1.611 | 0.107 |
| Mallya, Fiocco., (2019) | 0.400 | -0.124 | 0.924 | 1.496 | 0.135 |
| Zeidan et al., (2010) | 0.731 | 0.161 | 1.301 | 2.515 | 0.012* |
| Fixed | 0.385 | 0.115 | 0.656 | 2.791 | 0.005* |
| Random | 0.390 | 0.101 | 0.679 | 2.643 | 0.008* |

\*p<0.05 \*\*p<0.001

**Table 6.** Summary effect sizes of the Single-group studies (n = 2)

| Study name | Effect size | Lower limit | Upper limit | Z value | P value |
| --- | --- | --- | --- | --- | --- |
| (Bigelow et al., 2021) | 0.717 | 0.189 | 1.244 | 2.664 | 0.008* |
| (Xie et al., 2022) | 0.518 | 0.152 | 0.885 | 2.771 | 0.006* |
| Fixed | 0.583 | 0.282 | 0.884 | 3.796 | <0.001** |
| Random | 0.583 | 0.282 | 0.884 | 3.796 | <0.001** |

\*p<0.05 \*\*p<0.001

The heterogeneity test results for RCT two-group studies revealed a Q-value of 76.018, p-value < 0.001, and I-squared = 71.06. For CT two-group studies, the heterogeneity test showed a Q-value of 3.413, p-value = 0.332, and I-squared = 12.095. In single-group studies, the results were Q-value = 0.367, p-value = 0.545, and I-squared < 0.001. These findings suggest strong heterogeneity in RCT studies and weak heterogeneity in CT and single-group studies based on the Q-value test.

Additionally, the I-squared value confirms weak homogeneity in RCT studies and strong homogeneity in CT and single-group studies, aligning with the Q-value test results.

### Bias in meta-analysis results

Generally, a systematic review will have valid and high-quality results if efforts are made to minimize biases as much as possible. One such bias is language bias, where the researcher should attempt to include studies in another language. In this study, we included Persian articles in addition to English articles as an inclusion criterion, thereby reducing language bias in accordance with Cochrane Institute protocols. Another potential bias is selection bias, which occurs when the researcher’s judgment influences the selection. In this study, the use of a standardized protocol prevented the researcher’s intuition and judgment from influencing the selection of studies. The potential for publication bias to influence the results was also addressed using Duval and Tweedie’s trim and fill method and the Fail-Safe N test. We used JASP software (Love et al., 2019) to draw forest plots and funnel plots.

Sterne et al. (2011) suggest that a minimum of 10 studies is essential for constructing a reliable funnel plot. Since the present meta-analysis included an adequate number only of RCT studies exceeding this threshold, the evaluation of funnel plots was limited to these studies.

Egger regression analysis was performed to examine asymmetry in the funnel plot, which can indicate potential publication bias in the reviewed literature. Funnel plots are graphical representations of effect sizes plotted against their standard errors. Because larger sample sizes tend to provide more precise effect estimates (resulting in smaller standard errors), data points from smaller studies will show greater dispersion, while those from larger studies will be closer to the mean overall effect size. In the absence of publication bias, the distribution of these data points should be symmetrical around the mean effect size. However, bias is often evident when smaller studies with fewer adverse outcomes are reported, leading to asymmetry in the funnel plot. Egger regression quantitatively assesses the extent of this asymmetry (Egger et al., 1997).

According to Figures 5, funnel plots for the effects of mindfulness on working memory were calculated for RCTs. Egger’s regression test for asymmetry of funnel plots failed to reach statistical significance (z = 0.744, p = 0.457). In the Fail-Safe N test, findings are considered robust if (N ≥ 5n + 10), where (n) is the number of studies (Rosenberg, 2005). In this meta-analysis, the N value indicated that for our analysis of RCT study designs, 897 studies would have to be added to the suppression model to make our results not statistically significant. Therefore, the findings of our meta-analysis results for RCT studies are robust to possible publication bias.

**Figure 5.**
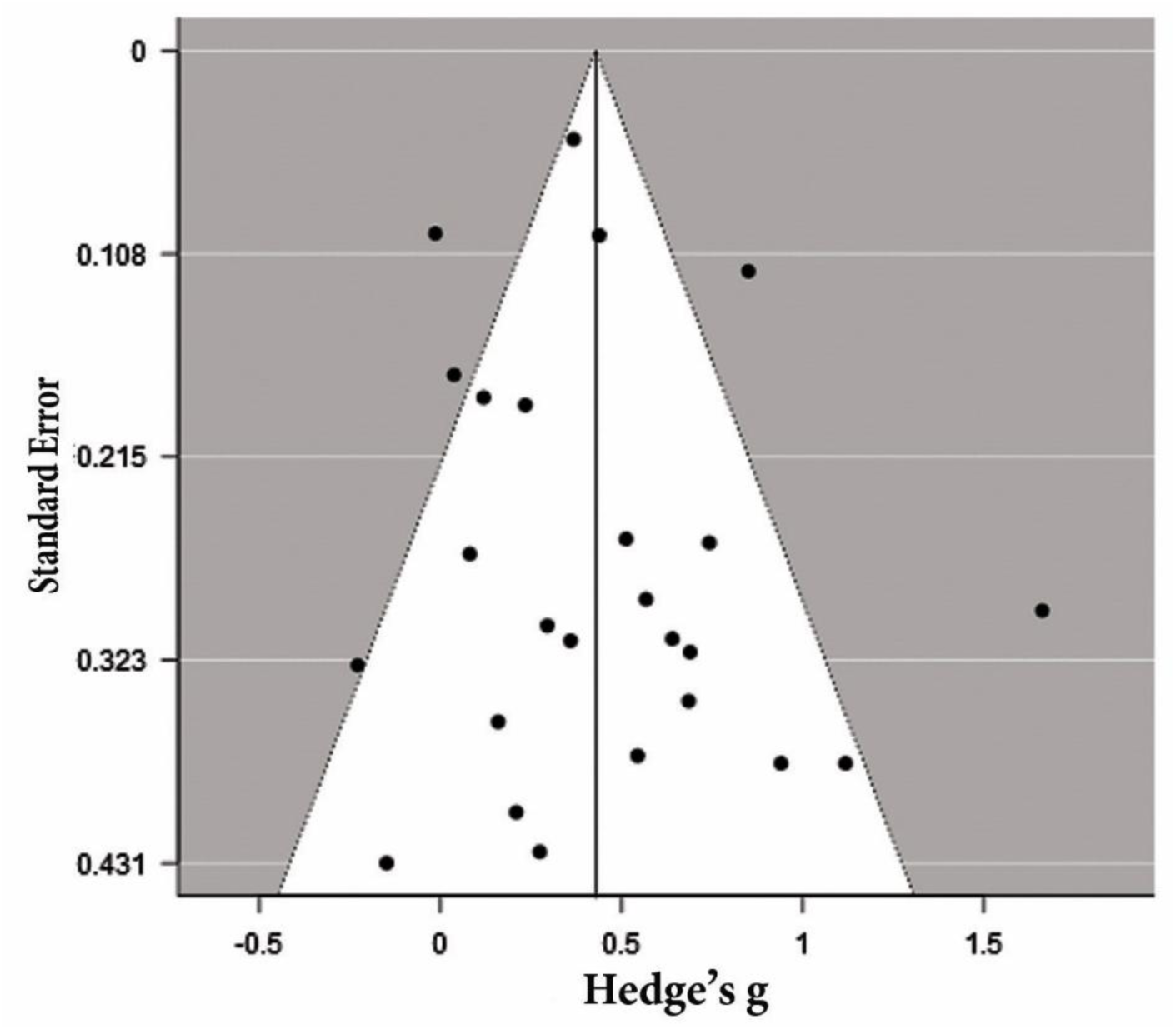
The funnel plot for the effects of mindfulness on working memory in RCT 2-group. Note: vertical line indicates overall effect size, circles indicate individual studies, and diagonal lines indicate 95% confidence intervals.

### Study Quality Assessment

Overall, the included studies were of medium quality, scoring an average of 3.75 out of 6 on the adapted version of the Delphi criteria. The majority of studies randomized participants to either the mindfulness or control condition (85% of included studies), defined eligibility criteria (85% of included studies), and provided sufficient statistics to enable effect size calculation for meta-analysis (100% of included studies). Most studies also reported that the mindfulness and control groups were demographically similar (65% of included studies). However, only a minority of studies applied an intention to treat analysis (or included all participants who provided baseline data in their analysis, 30% of studies), and very few included studies performed the assessment of working memory performance with the assessing researchers blinded to group allocation (10%). **Moderation Analyses**

The variables of control group type, clinical status, delivery format (FF/ON), duration of the intervention (DI), length of sessions (LS) and number of sessions (NS) were evaluated as moderators of the effects in the final meta-analysis dataset. When all potential moderator variables were included in the meta-analysis, the omnibus test of model coefficients was not significant: Q(6) = 0.890, p = 0.989. Additionally, no individual potential moderator approached significance (Table 7).

**Table 7.** Subgroup analyses of key moderators Moderator coefficients:

|  | Estimate | Standard Error | z | p | 95% Confidence Interval |  |
| --- | --- | --- | --- | --- | --- | --- |
|  |  |  |  |  | Lower | Upper |
| Control group | 0.110 | 0.254 | 0.432 | 0.665 | -0.388 | 0.607 |
| Clinical Status | 0.018 | 0.309 | 0.060 | 0.952 | -0.587 | 0.624 |
| NS | -0.107 | 0.372 | -0.288 | 0.774 | -0.836 | 0.622 |
| DI | -0.054 | 0.333 | -0.162 | 0.871 | -0.707 | 0.599 |
| LS | 0.136 | 0.264 | 0.513 | 0.608 | -0.382 | 0.654 |
| FF/ON | 0.131 | 0.279 | 0.468 | 0.640 | -0.416 | 0.677 |

FF: Face-to-face, ON: Online, DI: Duration of the intervention, NS: Number of Sessions, LS: Length of Sessions

## Discussion

The aim of the present study was to use meta-analysis to examine studies on the effects of mindfulness interventions on working memory in both healthy populations and patients with clinical disorders across different age groups. In this meta-analysis, 29 studies meeting the inclusion criteria were included, comprising a total of 2076 participants aged between 5 and 85 years. The current results showed a significant medium effect size (for RCT 2-group studies Hedges g = 0.438; for CT 2-group studies Hedges g = 0.385; for single-group studies Hedges g = 0.583). As such, our results indicate that mindfulness interventions have a positive effect on working memory. Our moderator analysis indicated that none of the moderating factors we examined had a significant effect, suggesting that the effect of mindfulness on working memory is not altered by factors like whether the intervention is taught online or face to face, whether short or long interventions are provided, or whether the intervention is provided to healthy or clinical populations.

Previous research has suggested that mindful awareness employs working memory to maintain awareness of moment-to-moment experiences (Geronimi et al., 2020). Meditation practice requires sustained attention while simultaneously directing attention to the present experience, closely related to working memory performance (Quach et al., 2016). As such, our results suggest that the working memory training implicit in mindfulness meditation practice may have a positive effect on working memory function across age groups and for both clinical and healthy populations.

Within this “working memory practice effect” explanation, the impact of mindfulness on working memory may be due to changes in levels of acceptance/non-judgmental action rather than due to attention training effects. The theoretical framework of mindfulness primarily relies on two factors: awareness and acceptance (Baer et al., 2006). However, it is unclear which factor of mindfulness might produce the effects we observed on working memory. Ruocco & Wonders, (2013) have suggested that the trait of mindfulness might not be related to working memory through increased awareness but rather through increased acceptance. The active process of acceptance can help maintain an individual’s attitude towards their experiences by relying on working memory capacities and actively using acceptance-based strategies (Levi & Rosenstreich, 2019). Conversely, Gallant (2016) has suggested that mindfulness meditation does not directly improve working memory; instead, the improvement is indirect, driven by a reduction in mind-wandering (Gallant, 2016). In alignment with this argument, it is possible that the impact of mind-wandering on working memory is greater under negative conditions. Interventions that prevent the increase of negative affect or alter the impact of mind-wandering on working memory can prevent working memory impairment (Banks et al., 2015). Irrespective of the answer to this question, the potential “indirect” nature of the effects of mindfulness on working memory does not negate the existence of the effect (Cásedas et al., 2020). Nonetheless, further research is required to better understand the mechanisms by which mindfulness improves working memory.

Alternatively, focused attention meditation may have an indirect effect on working memory capacity by reducing stress or increasing relaxation, thereby enhancing attention focus on current tasks (Jha et al., 2019). Studies have shown that mindfulness meditation protects working memory capacity and performance from stress-induced declines and negative impacts (Baranski & Was, 2018; Jha et al., 2017). Previous studies on individual differences in working memory capacity and emotion regulation report that working memory capacity is not related to the natural expression or experience of emotions but rather to the ability to successfully regulate emotions (Schmeichel et al., 2008; Schmeichel & Demaree, 2010). Studies indicate that the impact of mindfulness on working memory is associated with a reduction in negative affect rather than positive affect. It appears that mindfulness improves working memory more through regulating negative emotions because only the experience of negative affect requires regulation (Jha et al., 2010). Mechanistically, the non-judgmental observation fostered by mindfulness may neutralize the negative effect of a stimulus. In other words, mindfulness may reduce the explicit processing of the negative content of a stimulus, thereby reducing its impact on memory. As such, mindful awareness might be a way to reduce the impact of negative events on memory (Alberts & Thewissen, 2011). Finally, it is possible that the mechanisms underlying the effects of mindfulness on working memory that we have observed are a combination of these two mechanisms, with the relative contribution of each mechanistic facet varying across individuals.

Despite the positive results in our meta-analysis, we note that the present findings contradict some previous studies. This may be due to differences in methodological approaches for calculating effect sizes or differences in inclusion criteria (Im et al., 2021). For instance, Im et al. (2021) conducted a meta-analysis that focused on six studies involving participants aged 18 to 59 who presented with psychiatric or medical conditions. The cognitive assessment tasks employed in their analysis included the Digit Span, Letter Number Sequencing, and Operation Span Task. In a separate systematic review and meta-analysis, Millett et al. (2021) conducted a review of nine papers to examine the impact of mindfulness programs on executive performance in a sample of healthy individuals aged 18 years and older. Similarly, Yakobi et al. (2021) selected healthy adults from eight articles for their meta-analysis, with participants ranging in age from 18 to 65 years. These differences are noteworthy considering that the present study included both clinical and non-clinical subjects across all age groups. Additionally, all articles featuring any form of cognitive task that evaluated working memory were included in the analysis, resulting in a total of 29 articles focused on working memory. Although the current study suggests that mindfulness has a positive effect on working memory function, further research is needed to address methodological challenges in meta-analysis and the limitations of existing studies to reach a definitive conclusion. As mentioned, the current meta-analysis combines data from studies on MBIs conducted with diverse populations and provides an estimate of the broad effects of MBIs on working memory. However, we identified several methodological issues in the studies, which warrant highlighting to enable recommendations for further research.

First, we noticed significant differences in the conceptualizations and operational definitions of mindfulness. The studies included in this research used various types of mindfulness interventions, with different numbers of sessions (ranging from one to 30 sessions), varying durations of practice (from 5 minutes to 3 hours), and mindfulness instructors with different levels of experience. Additionally, a number of the studies we included provided mindfulness sessions in an online format, in contrast to the typical in-person format, which could have different impacts on participants. However, our moderator analysis suggested that the effects did not differ between these types of studies. While the diversity among MBIs is a current limitation, our results suggest that future reviews might encompass a broader range of interventions. There is an increasing interest in using digital and web-based programs, and our results suggest the effects of these programs on working memory do not differ from traditional face-to-face approaches. We also note that the positive results we detected in the context of the broad variations between studies might be viewed as a strength of our study and the literature on the effects of mindfulness on working memory – indicating that our conclusions likely generalize across different mindfulness program types, durations, instructor experiences and populations.

Second, most reviewed studies did not indicate whether participants had prior experience with mindfulness practices, nor report the participants’ level of interest and engagement. Variability in participants’ motivation and adherence, and its effects in MBI studies as a moderator, is relatively under-researched, and less than half of the studies included here reported adherence and motivation data. This prevented us from assessing adherence as a hypothetical moderator. Recent theoretical explanations for the limited effects of mindfulness in schools have suggested that lack of adherence to home practice recommendations may be a critical factor in null results (Strohmaier & Bailey, 2023), so future research should explore adherence to home practice recommendations in more detail. Most studies also did not test whether mindfulness skills improved after receiving MBIs, and the follow-up period was not examined in many studies. Since putative mechanistic pathways for the effects of mindfulness on working memory that we observed is that MBIs increase mindfulness skills, which in turn leads to improved working memory performance, demonstrating that the MBIs do indeed increase mindfulness skills prior to the working memory changes in future research is important. Without such information, the internal validity of the effects of an MBI program on working memory is reduced, and significant findings cannot be solely attributed to the treatment effect. Furthermore, variation in participants’ engagement with mindfulness practices post-intervention may also contribute to long-term cognitive effects. Assessing this requires follow-up evaluations post-intervention, which were generally lacking among the present studies. If possible, including follow-up time points in future MBI trials will help clarify the potential role of continued mindfulness practice on working memory function.

Third, sample sizes in the studies varied substantially (from 16 to 307 participants). 13 studies had small sample sizes, with a total of 40 participants or fewer, and only four studies used recommended sample size determination software before commencing their study (Faul et al., 2009). In this context, we also note that when multiple dependent variables are used, the estimated sample size should be increased, a point that future research should account for. Studies with small sample sizes significantly reduce reproducibility due to insufficient power. Therefore, we recommend that future studies conduct power analysis and use appropriate sample sizes through sample size determination software.

Fourth, the studies included varying proportions of genders, and some studies included only one gender (male). Given that gender can influence the outcomes of mindfulness interventions (Wang & Chopel, 2017), future research might be recommended to determine whether the effects of mindfulness on working memory vary with gender. Fifth, the results of our study should be interpreted with caution because participants in the included studies had different age ranges (from 5 to 85 years), and in nine studies, adolescent, young adult, and elderly ages were combined. Since empirical evidence suggests that older participants show greater effects from MBIs than younger participants (Alispahic & Hasanbegovic-Anic, 2017), this may explain the inconsistent findings in studies using participants with different age ranges. Therefore, more research is needed to systematically evaluate the impact of MBIs on different age groups.

The sixth limitation is the lack of active control groups in some studies to account for the effects of confounding factors and controls such as non-specific intervention effects, participant characteristics, group differences, and motivation. When groups are not matched based on the above factors, the internal validity of study results is weakened, which may risk assessment bias. Although our analysis did not show differences depending on whether studies included an active or non-active control group, future research should consider an active control group with programs like cognitive enhancement programs that control for confounding variables (Tang et al., 2015). The aforementioned limitations necessitate caution in interpreting the present findings as potential confounding factors and unexplained error variance, which may have biased the results of individual studies and consequently affected the current meta-analysis results. Therefore, there is a clear need for methodologically robust randomized controlled trials with large sample sizes that can establish a clear link between mindfulness components and working memory. It is also suggested that MBI studies be conducted with the integration of neurocognitive measures. In this meta-analysis, five studies used this method. Since electroencephalography (EEG) is suitable for studying time-sensitive executive function processes (Falkenstein et al., 1999; Van Veen & Carter, 2002), it is suggested to use this method to examine changes in working memory, as existing research indicates that neural activity related to working memory differs in experienced meditators (Bailey et al., 2020; Wang et al., 2020). Finally, due to the theorized relationship between mindfulness components and specific neurocognitive systems, this study examined the development of mindfulness capacity in relation to improving working memory (Vago & Silbersweig, 2012). However, assessing whether cognitive improvements truly parallel increased mindfulness capacity was beyond the scope of this review. Future researchers are encouraged to investigate these interesting and complex relationships.

## References

1. Alberts, H. J., & Thewissen, R. (2011). The effect of a brief mindfulness intervention on memory for positively and negatively valenced stimuli. Mindfulness, 2, 73–77.

2. Alispahic, S., & Hasanbegovic-Anic, E. (2017). Mindfulness: Age and gender differences on a Bosnian sample. Psychological Thought, 10(1), 155–166.

3. Alizadehgoradel, J., Imani, S., Nejati, V., & Fathabadi, J. (2019). Mindfulness-based substance abuse treatment (MBSAT) improves executive functions in adolescents with substance use disorders. *Neurology*, Psychiatry and Brain Research, 34, 13–21.

4. Axelsen, J. L., Meline, J. S. J., Staiano, W., & Kirk, U. (2022). Mindfulness and music interventions in the workplace: Assessment of sustained attention and working memory using a crowdsourcing approach. BMC Psychology, 10(1), 108.

5. Bachmann, K., Lam, A. P., Sörös, P., Kanat, M., Hoxhaj, E., Matthies, S., Feige, B., Müller, H., Özyurt, J., & Thiel, C. M. (2018). Effects of mindfulness and psychoeducation on working memory in adult ADHD: A randomised, controlled fMRI study. Behaviour Research and Therapy, 106, 47–56.

6. Baer, R. A., Smith, G. T., Hopkins, J., Krietemeyer, J., & Toney, L. (2006). Using self-report assessment methods to explore facets of mindfulness. Assessment, 13(1), 27–45.

7. Bailey, N. W., Freedman, G., Raj, K., Spierings, K. N., Piccoli, L. R., Sullivan, C. M., Chung, S. W., Hill, A. T., Rogasch, N. C., & Fitzgerald, P. B. (2020). Mindfulness meditators show enhanced accuracy and different neural activity during working memory. Mindfulness, 11, 1762–1781.

8. Banks, J. B., Welhaf, M. S., & Srour, A. (2015). The protective effects of brief mindfulness meditation training. Consciousness and Cognition, 33, 277–285.

9. Baranski, M. F., & Was, C. A. (2018). A more rigorous examination of the effects of mindfulness meditation on working memory capacity. Journal of Cognitive Enhancement, 2, 225–239.

10. Basso, J. C., McHale, A., Ende, V., Oberlin, D. J., & Suzuki, W. A. (2019). Brief, daily meditation enhances attention, memory, mood, and emotional regulation in non-experienced meditators. Behavioural Brain Research, 356, 208–220.

11. Bigelow, H., Gottlieb, M. D., Ogrodnik, M., Graham, J. D., & Fenesi, B. (2021). The differential impact of acute exercise and mindfulness meditation on executive functioning and psycho-emotional well-being in children and youth with ADHD. Frontiers in Psychology, 12, 660845.

12. Brisbon, N. M., & Lachman, M. E. (2017). Dispositional mindfulness and memory problems: The role of perceived stress and sleep quality. Mindfulness, 8, 379–386.

13. Brown, K. W., Goodman, R. J., Ryan, R. M., & Anālayo, B. (2016). Mindfulness enhances episodic memory performance: Evidence from a multimethod investigation. PloS One, 11(4), e0153309.

14. Brown, K. W., & Ryan, R. M. (2003). The benefits of being present: Mindfulness and its role in psychological well-being. Journal of Personality and Social Psychology, 84(4), 822.

15. Cásedas, L., Pirruccio, V., Vadillo, M. A., & Lupiáñez, J. (2020). Does mindfulness meditation training enhance executive control? A systematic review and meta-analysis of randomized controlled trials in adults. Mindfulness, 11, 411–424.

16. Chambers, R., Lo, B. C. Y., & Allen, N. B. (2008). The impact of intensive mindfulness training on attentional control, cognitive style, and affect. Cognitive Therapy and Research, 32, 303–322.

17. Chiesa, A., Calati, R., & Serretti, A. (2011). Does mindfulness training improve cognitive abilities? A systematic review of neuropsychological findings. Clinical Psychology Review, 31(3), 449–464.

18. Dai, M., Li, Y., Gan, S., & Du, F. (2019). The reliability of estimating visual working memory capacity. Scientific Reports, 9(1), 1155.

19. De Grijs, R., Wicker, J. E., & Bono, G. (2014). Clustering of local group distances: Publication bias or correlated measurements? I. The Large Magellanic Cloud. The Astronomical Journal, 147(5), 122.

20. Dubert, C. J., Schumacher, A. M., Locker, L., Gutierrez, A. P., & Barnes, V. A. (2016). Mindfulness and emotion regulation among nursing students: Investigating the mediation effect of working memory capacity. Mindfulness, 7, 1061–1070.

21. Duval, S., & Tweedie, R. (2000). Trim and fill: A simple funnel-plot–based method of testing and adjusting for publication bias in meta-analysis. Biometrics, 56(2), 455–463.

22. Egger, M., Smith, G. D., Schneider, M., & Minder, C. (1997). Bias in meta-analysis detected by a simple, graphical test. Bmj, 315(7109), 629–634.

23. Eisenbeck, N., Luciano, C., & Valdivia-Salas, S. (2018). Effects of a focused breathing mindfulness exercise on attention, memory, and mood: The importance of task characteristics. Behaviour Change, 35(1), 54–70.

24. Falkenstein, M., Hoormann, J., & Hohnsbein, J. (1999). ERP components in Go/Nogo tasks and their relation to inhibition. Acta Psychologica, 101(2–3), 267–291.

25. Faul, F., Erdfelder, E., Buchner, A., & Lang, A.-G. (2009). Statistical power analyses using G* Power 3.1: Tests for correlation and regression analyses. Behavior Research Methods, 41(4), 1149–1160.

26. Flook, L., Hirshberg, M. J., Gustafson, L., McGehee, C., Knoeppel, C., Tello, L. Y., Bolt, D. M., & Davidson, R. J. (2024). Mindfulness training enhances students’ executive functioning and social emotional skills. Applied Developmental Science, 1–20.

27. Gallant, S. N. (2016). Mindfulness meditation practice and executive functioning: Breaking down the benefit. Consciousness and Cognition, 40, 116–130.

28. Geronimi, E. M., Arellano, B., & Woodruff-Borden, J. (2020). Relating mindfulness and executive function in children. Clinical Child Psychology and Psychiatry, 25(2), 435–445.

29. Ghorbani, M., & Khalilian, R. (2016). Effects of Mindfulness Training on Working Memory and Behavioral Inhibition for Adults with Attention Deficit / Hyperactivity. Icss, 18(3), 90–100.

30. Gill, L.-N., Renault, R., Campbell, E., Rainville, P., & Khoury, B. (2020). Mindfulness induction and cognition: A systematic review and meta-analysis. Consciousness and Cognition, 84, 102991.

31. Goyal, M., Singh, S., Sibinga, E. M., Gould, N. F., Rowland-Seymour, A., Sharma, R., Berger, Z., Sleicher, D., Maron, D. D., & Shihab, H. M. (2014). Meditation programs for psychological stress and well-being: A systematic review and meta-analysis. JAMA Internal Medicine, 174(3), 357–368.

32. Greenberg, J., Romero, V. L., Elkin-Frankston, S., Bezdek, M. A., Schumacher, E. H., & Lazar, S. W. (2019). Reduced interference in working memory following mindfulness training is associated with increases in hippocampal volume. Brain Imaging and Behavior, 13, 366–376.

33. Hedge, C., Powell, G., & Sumner, P. (2018). The reliability paradox: Why robust cognitive tasks do not produce reliable individual differences. Behavior Research Methods, 50, 1166–1186.

34. Hedges, L. V. (1981). Distribution theory for Glass’s estimator of effect size and related estimators. Journal of Educational Statistics, 6(2), 107–128.

35. Hedges, L. V., & Olkin, I. (2014). Statistical methods for meta-analysis. Academic press.

36. Higgins, J. P., & Thompson, S. G. (2002). Quantifying heterogeneity in a meta-analysis. Statistics in Medicine, 21(11), 1539–1558.

37. Higgins, J. P., Thompson, S. G., Deeks, J. J., & Altman, D. G. (2003). Measuring inconsistency in meta-analyses. Bmj, 327(7414), 557–560.

38. Huedo-Medina, T. B., Sánchez-Meca, J., Marín-Martínez, F., & Botella, J. (2006). Assessing heterogeneity in meta-analysis: Q statistic or I^2^ index? Psychological Methods, 11(2), 193.

39. Im, S., Stavas, J., Lee, J., Mir, Z., Hazlett-Stevens, H., & Caplovitz, G. (2021). Does mindfulness-based intervention improve cognitive function?: A meta-analysis of controlled studies. Clinical Psychology Review, 84, 101972.

40. Isham, A. E., del Palacio-Gonzalez, A., & Dritschel, B. (2020). Trait mindfulness and emotion regulation upon autobiographical memory retrieval during depression remission. Mindfulness, 11(12), 2828– 2840.

41. Jansen, P., Dahmen-Zimmer, K., Kudielka, B. M., & Schulz, A. (2017). Effects of karate training versus mindfulness training on emotional well-being and cognitive performance in later life. Research on Aging, 39(10), 1118–1144.

42. Jensen, C. G., Vangkilde, S., Frokjaer, V., & Hasselbalch, S. G. (2012). Mindfulness training affects attention—Or is it attentional effort? Journal of Experimental Psychology: General, 141(1), 106.

43. Jha, A. P., Denkova, E., Zanesco, A. P., Witkin, J. E., Rooks, J., & Rogers, S. L. (2019). Does mindfulness training help working memory ‘work’better? Current Opinion in Psychology, 28, 273–278.

44. Jha, A. P., Stanley, E. A., Kiyonaga, A., Wong, L., & Gelfand, L. (2010). Examining the protective effects of mindfulness training on working memory capacity and affective experience. Emotion, 10(1), 54.

45. Jha, A. P., Witkin, J. E., Morrison, A. B., Rostrup, N., & Stanley, E. (2017). Short-form mindfulness training protects against working memory degradation over high-demand intervals. Journal of Cognitive Enhancement, 1, 154–171.

46. Jha, A. P., Zanesco, A. P., Denkova, E., Rooks, J., Morrison, A. B., & Stanley, E. A. (2020). Comparing mindfulness and positivity trainings in high-demand cohorts. Cognitive Therapy and Research, 44, 311–326.

47. Kabat-Zinn, J. (1990). Full Catastrophe Living: Using the Wisdom of Your Body and Mind to Face Stress, Pain, and Illness. New York: Delacorte Press.

48. Kiani, B., Hadianfard, H., & Mitchell, J. T. (2017). The impact of mindfulness meditation training on executive functions and emotion dysregulation in an Iranian sample of female adolescents with elevated attention-deficit/hyperactivity disorder symptoms. Australian Journal of Psychology, 69(4), 273–282.

49. Lakens, D. (2013). Calculating and reporting effect sizes to facilitate cumulative science: A practical primer for t-tests and ANOVAs. Frontiers in Psychology, 4, 863.

50. Lao, S.-A., Kissane, D., & Meadows, G. (2016). Cognitive effects of MBSR/MBCT: A systematic review of neuropsychological outcomes. Consciousness and Cognition, 45, 109–123.

51. Lebares, C. C., Guvva, E. V., Olaru, M., Sugrue, L. P., Staffaroni, A. M., Delucchi, K. L., Kramer, J. H., Ascher, N. L., & Harris, H. W. (2019). Efficacy of mindfulness-based cognitive training in surgery: Additional analysis of the mindful surgeon pilot randomized clinical trial. JAMA Network Open, 2(5), e194108–e194108.

52. Levi, U., & Rosenstreich, E. (2019). Mindfulness and memory: A review of findings and a potential model. Journal of Cognitive Enhancement, 3(3), 302–314.

53. Love, J., Selker, R., Marsman, M., Jamil, T., Dropmann, D., Verhagen, J., Ly, A., Gronau, Q. F., Šmíra, M., & Epskamp, S. (2019). JASP: Graphical statistical software for common statistical designs. Journal of Statistical Software, 88, 1–17.

54. Lovette, B. C., Kanaya, M. R., Bannon, S. M., Vranceanu, A.-M., & Greenberg, J. (2022). “Hidden gains”? Measuring the impact of mindfulness-based interventions for people with mild traumatic brain injury: A scoping review. Brain Injury, 36(9), 1059–1070.

55. Lueke, A., & Lueke, N. (2019). Mindfulness improves verbal learning and memory through enhanced encoding. Memory & Cognition, 47(8), 1531–1545.

56. Mak, C., Whittingham, K., Cunnington, R., & Boyd, R. N. (2018). Efficacy of mindfulness-based interventions for attention and executive function in children and adolescents—A systematic review. Mindfulness, 9, 59–78.

57. Makmee, P. (2022). Increasing attention and working memory in elementary students using mindfulness training programs. FWU Journal of Social Sciences, 16(3), 107–119.

58. Melby-Lervåg, M., & Hulme, C. (2013). Is working memory training effective? A meta-analytic review. Developmental Psychology, 49(2), 270.

59. Millett, G., D’Amico, D., Amestoy, M. E., Gryspeerdt, C., & Fiocco, A. J. (2021a). Do group-based mindfulness meditation programs enhance executive functioning? A systematic review and meta-analysis of the evidence. Consciousness and Cognition, 95, 103195.

60. Millett, G., D’Amico, D., Amestoy, M. E., Gryspeerdt, C., & Fiocco, A. J. (2021b). Do group-based mindfulness meditation programs enhance executive functioning? A systematic review and meta-analysis of the evidence. Consciousness and Cognition, 95, 103195.

61. Mirabito, G., & Verhaeghen, P. (2023). The effects of mindfulness interventions on older adults’ cognition: A Meta-analysis. The Journals of Gerontology: Series B, 78(3), 394–408.

62. Mitchell, J. T., McIntyre, E. M., English, J. S., Dennis, M. F., Beckham, J. C., & Kollins, S. H. (2017). A pilot trial of mindfulness meditation training for ADHD in adulthood: Impact on core symptoms, executive functioning, and emotion dysregulation. Journal of Attention Disorders, 21(13), 1105– 1120.

63. Moher, D., Liberati, A., Tetzlaff, J., Altman, D. G., & PRISMA Group*, t. (2009). Preferred reporting items for systematic reviews and meta-analyses: The PRISMA statement. Annals of Internal Medicine, 151(4), 264–269.

64. Morrison, A. B., Goolsarran, M., Rogers, S. L., & Jha, A. P. (2014). Taming a wandering attention: Short-form mindfulness training in student cohorts. Frontiers in Human Neuroscience, 7, 897.

65. Morrison, A. B., & Jha, A. P. (2015). Mindfulness, attention, and working memory. Handbook of Mindfulness and Self-Regulation, 33–45.

66. Mrazek, M. D., Franklin, M. S., Phillips, D. T., Baird, B., & Schooler, J. W. (2013). Mindfulness training improves working memory capacity and GRE performance while reducing mind wandering. Psychological Science, 24(5), 776–781.

67. Nakagawa, S., Lagisz, M., Jennions, M. D., Koricheva, J., Noble, D. W., Parker, T. H., Sánchez-Tójar, A., Yang, Y., & O’Dea, R. E. (2022). Methods for testing publication bias in ecological and evolutionary meta-analyses. Methods in Ecology and Evolution, 13(1), 4–21.

68. Orwin, R. G. (1983). A fail-safe N for effect size in meta-analysis. Journal of Educational Statistics, 8(2), 157–159.

69. Podina, I. R., Mogoase, C., David, D., Szentagotai, A., & Dobrean, A. (2016). A meta-analysis on the efficacy of technology mediated CBT for anxious children and adolescents. Journal of Rational-Emotive & Cognitive-Behavior Therapy, 34, 31–50.

70. Quach, D., Mano, K. E. J., & Alexander, K. (2016). A randomized controlled trial examining the effect of mindfulness meditation on working memory capacity in adolescents. Journal of Adolescent Health, 58(5), 489–496.

71. Quek, F. Y., Majeed, N. M., Kothari, M., Lua, V. Y., Ong, H. S., & Hartanto, A. (2021). Brief mindfulness breathing exercises and working memory capacity: Findings from two experimental approaches. Brain Sciences, 11(2), 175.

72. Roeser, R. W., Schonert-Reichl, K. A., Jha, A., Cullen, M., Wallace, L., Wilensky, R., Oberle, E., Thomson, K., Taylor, C., & Harrison, J. (2013). Mindfulness training and reductions in teacher stress and burnout: Results from two randomized, waitlist-control field trials. Journal of Educational Psychology, 105(3), 787.

73. Rosenberg, M. S. (2005). The file-drawer problem revisited: A general weighted method for calculating fail-safe numbers in meta-analysis. Evolution, 59(2), 464–468.

74. Rosenthal, R. (1979). The file drawer problem and tolerance for null results. Psychological Bulletin, 86(3), 638.

75. Ruocco, A. C., & Wonders, E. (2013). Delineating the contributions of sustained attention and working memory to individual differences in mindfulness. Personality and Individual Differences, 54(2), 226–230.

76. Scharfen, J., Jansen, K., & Holling, H. (2018). Retest effects in working memory capacity tests: A meta-analysis. Psychonomic Bulletin & Review, 25(6), 2175–2199.

77. Schmeichel, B. J., & Demaree, H. A. (2010). Working memory capacity and spontaneous emotion regulation: High capacity predicts self-enhancement in response to negative feedback. Emotion, 10(5), 739.

78. Schmeichel, B. J., Volokhov, R. N., & Demaree, H. A. (2008). Working memory capacity and the self-regulation of emotional expression and experience. Journal of Personality and Social Psychology, 95(6), 1526.

79. Shemesh, L., Mendelsohn, A., Panitz, D. Y., & Berkovich-Ohana, A. (2023). Enhanced declarative memory in long-term mindfulness practitioners. Psychological Research, 87(1), 294–307.

80. Sterne, J. A., Sutton, A. J., Ioannidis, J. P., Terrin, N., Jones, D. R., Lau, J., Carpenter, J., Rücker, G., Harbord, R. M., & Schmid, C. H. (2011). Recommendations for examining and interpreting funnel plot asymmetry in meta-analyses of randomised controlled trials. Bmj, 343.

81. Strohmaier, S., & Bailey, N. W. (2023). Do not keep calm and carry on: School-based mindfulness programmes should test making mindfulness practice available in the school day. Mindfulness, 14(12), 3086–3097.

82. Tang, Y.-Y., Hölzel, B. K., & Posner, M. I. (2015). The neuroscience of mindfulness meditation. Nature Reviews Neuroscience, 16(4), 213–225.

83. Uopasai, S., Bunterm, T., Tang, K. N., & Saksangawong, C. (2022). The effect of meditation on metacognitive ability, working memory ability, academic achievement, and stress levels. *Humanities, Arts and Social Sciences Studies (Former Name Silpakorn University Journal of Social Sciences*, Humanities, and Arts), 217–226.

84. Vago, D. R., & Silbersweig, D. A. (2012). Self-awareness, self-regulation, and self-transcendence (S-ART): A framework for understanding the neurobiological mechanisms of mindfulness. Frontiers in Human Neuroscience, 6, 296.

85. Van Veen, V., & Carter, C. S. (2002). The anterior cingulate as a conflict monitor: fMRI and ERP studies. Physiology & Behavior, 77(4–5), 477–482.

86. Verhagen, A. P., De Vet, H. C., De Bie, R. A., Kessels, A. G., Boers, M., Bouter, L. M., & Knipschild, P. G. (1998). The Delphi list: A criteria list for quality assessment of randomized clinical trials for conducting systematic reviews developed by Delphi consensus. Journal of Clinical Epidemiology, 51(12), 1235–1241.

87. Vieth, E., & von Stockhausen, L. (2022). Mechanisms underlying cognitive effects of inducing a mindful state. Journal of Cognition, 5(1).

88. Wang, M. Y., Freedman, G., Raj, K., Fitzgibbon, B. M., Sullivan, C., Tan, W.-L., Van Dam, N., Fitzgerald, P. B., & Bailey, N. W. (2020). Mindfulness meditation alters neural activity underpinning working memory during tactile distraction. *Cognitive, Affective*, & Behavioral Neuroscience, 20, 1216– 1233.

89. Wang, W., & Chopel, T. (2017). Mindfulness and gender: A pilot quantitative study. Issues in Information Systems, 18(4).

90. Whitfield, T., Barnhofer, T., Acabchuk, R., Cohen, A., Lee, M., Schlosser, M., Arenaza-Urquijo, E. M., Böttcher, A., Britton, W., & Coll-Padros, N. (2022). The effect of mindfulness-based programs on cognitive function in adults: A systematic review and meta-analysis. Neuropsychology Review, 32(3), 677–702.

91. Xie, S., Gong, C., Lu, J., Li, H., Wu, D., Chi, X., & Chang, C. (2022). Enhancing Chinese preschoolers’ executive function via mindfulness training: An fNIRS study. Frontiers in Behavioral Neuroscience, 16, 961797.

92. Yakobi, O., Smilek, D., & Danckert, J. (2021). The effects of mindfulness meditation on attention, executive control and working memory in healthy adults: A meta-analysis of randomized controlled trials. Cognitive Therapy and Research, 1–18.

93. Yamaya, N., Tsuchiya, K., Takizawa, I., Shimoda, K., Kitazawa, K., & Tozato, F. (2021). Effect of one-session focused attention meditation on the working memory capacity of meditation novices: A functional near-infrared spectroscopy study. Brain and Behavior, 11(8), e2288.

94. Zainal, N. H., & Newman, M. G. (2024). Mindfulness enhances cognitive functioning: A meta-analysis of 111 randomized controlled trials. Health Psychology Review, 18(2), 369–395.

95. Zeidan, F., Johnson, S. K., Diamond, B. J., David, Z., & Goolkasian, P. (2010). Mindfulness meditation improves cognition: Evidence of brief mental training. Consciousness and Cognition, 19(2), 597– 605.

